# High G-Forces in Unintentionally Improper Infant Handling: Implications for Shaken Baby Syndrome Diagnosis

**DOI:** 10.1101/2024.10.31.621215

**Authors:** Jonathan S. Lee-Confer, Lila T. Wayman, Kathryn L. Havens

## Abstract

**Introduction:** Abusive head trauma (AHT), commonly referred to as Shaken Baby Syndrome (SBS), is diagnosed in approximately 33 per 100,000 infants annually in the United States. Traditional diagnostic criteria for SBS include subdural, subarachnoid, and retinal hemorrhages. While intentional shaking is a known cause, the potential for similar forces acting on the head resulting from accidental trauma has not been fully explored. This study investigates the biomechanical forces on a model infant’s head during improper handling to determine if such forces could contribute to SBS without malicious intent.

**Methods:** A realistic silicone infant model was equipped with an inertial measurement unit (IMU) to quantify head accelerations during two conditions: (1) placement of the infant model on a table with the head unsupported, and (2) manual shaking at maximum effort by 2 participants holding the model by the torso. Peak head accelerations were recorded for both conditions, and the results were analyzed for comparative assessment of the forces involved.

**Results:** The average peak head acceleration when placing the infant model on a table with the head unsupported was +30,952.67 ± 6,540.79 mg, with a range of +19,234.40 to +43,406.30 mg across trials. The average peak head accelerations during maximum effort shaking were significantly lower than placing the infant on the table, averaging 11,430.48 ± 9,539.06,867 vs. 30,952.67 ± 6,540.79 mg, p < 0.0001). There were no significant differences in head accelerations between participants when placing the infant model on the table with the head unsupported (p = 0.93) nor with shaking the baby with maximum effort (p = 0.97).

**Discussion:** The G-force in this study resulted in higher forces than experienced in an 18-mph car crash and a 5-mph bumper car collision. The study highlights that even accidental non-recommended handling of infants can result in high G-forces to the head, potentially leading to injuries similar to those observed in SBS. These findings highlight the necessity of supporting an infant’s head during handling and warrants caution against prematurely attributing physical abuse in SBS cases without considering unintentional causes.

## Introduction

Annually, approximately 33 per 100,000 are diagnosed with abusive head trauma (AHT), also known as Shaken baby syndrome (SBS), in hospitals within the United States [1]. Infant AHT generally refers to conditions resulting with or without direct impact to the skull which leads to injuries within the cranium [2]. The diagnosis of SBS is often based on signs such as subdural, subarachnoid, and retinal hemorrhages [3]. While intentional shaking is widely recognized as a cause of these injuries, the potential for similar forces resulting from improper or accidental handling of infants has not been fully explored.

Contrary to the implications of the phrase “Shaken Baby Syndrome,” evidence shows that the act of shaking alone may not generate sufficient forces to cause the severe intracranial injuries typically associated with SBS. These studies involve infant models to prevent harm to an actual infant, and have reported that head accelerations during shaking generally peak around 8.5 G-force [4]. In a another study, similar results of shaking a baby produced head accelerations of 10-15 G-forces [5] and another producing mean peak head accelerations of 18.6 G-forces in the fore-aft direction [6].

Because there is not a validated threshold of G-force that definitively cause infant head injuries, comparing magnitude of forces during different tasks allows for inference of potential trauma. Car safety experiments demonstrate 29.3 G-forces with simulated 18-mph car accidents and 5.2 G-forces in 5-mph bumper car collisions. These simulated collisions have adult dummies in car seats with the body in a semi-reclined position. This is similar to the position of infants during the head shaking tasks described above. However, other researchers have shown that impacts from babies falling over 1 meter falls result in head accelerations over 200 G-force [7]. This much larger force may be generated both because of the supine position of the body, and the direct impact of the head with a surface.

While research has focused on head accelerations resulting from shaking or falls, less attention has been given to other scenarios such as placing infants on hard surfaces without proper head support. The potential for such actions to produce high G-forces and cause trauma comparable to SBS remains poorly understood, especially in cases of accidental abusive head trauma. As such, the purpose of this study was to determine infant head G-forces in two conditions: 1) shaking a baby at maximum force, and 2) placing the infant on a table with the head unsupported. To ensure repeatability, we compared forces between two participants, and to identify whether differences in forces exist between placing on a table and forces identified in other studies during shaking, we compared forces during these tasks. We hypothesized that head accelerations would be significantly higher in placing an infant on a table with the head unsupported compared to shaking an infant at maximum force.

## Methods

### Case Review

#### Incident

A 34-week prematurely born 8-month-old female infant was brought to local hospital in Texas after a call was made to 911 from a distressed father who stated his baby became non-responsive and stopped breathing. The infant was 64.0cm in length and 8.62 kg in weight. The father brought his baby into the bedroom, placed her on the bed while he used the restroom briefly and then changed his shirt. Upon changing shirts and looking at his baby, the infant’s shirt was wet and covered in her own spit. The father got baby wipes on the other side of the room, walked back to the bed and the baby was making gurgling sounds. The father picked up the infant, turned her into a prone position, supported the infant’s body with one hand leaving the head unsupported and delivered back blows to get out whatever she choked on. The infant jerked and went limp. The father called 911 and ran out to the kitchen holding his infant. The infant was placed on the kitchen table, potentially with the head unsupported, and 911 was held on speaker phone. After some time, the father picked up the infant rapidly, ran to the front door and placed the infant on the grass while waiting for the ambulance to arrive. The infant did not survive.

#### Medical Records

The autopsy reports reported diagnoses of: blunt force injuries of the head and neck and bilateral panlobar acute pneumonia. Specifically, there were diagnoses of bilateral subdural hemorrhage (2 mL total), bilateral subarachnoid hemorrhage, intraventricular hemorrhage, bilateral optic nerve hemorrhages, COVID-19 positive, and posterior C1-C2 intramuscular hemorrhages without associated fractures. For the infant’s body measurements respective to age, the body length was less than 1^st^ percentile, weight 70^th^ percentile, head circumference of 43.0 cm in the 23^rd^ percentile, chest circumference of 43.0 cm in the 28^th^ percentile and abdominal circumference of 42.0 cm. The conclusion of the medical report is that this was a case of infant homicide. The infant born premature had early respiratory issues and the pneumonia may have been related to the positive COVID-19 diagnosis.

#### Instrumentation

One silicone life-like infant model was purchased for testing purposes. The height of the infant was 55.88 cm and the weight of the infant was 5.08 kg. This was slightly shorter and quite a bit lighter than the infant in the case,

The data collection was conducted in the SensorLab at the University of Arizona. Three-dimensional motion analysis of the infant’s head was conducted with one wireless Ultium inertial measurement unit (IMU) system collecting at 200 Hz (Noraxon, Scottsdale, AZ, USA). The IMU was securely affixed to the model infant’s head using a Velcro strap with the IMU at the anterior cranial just above the eyes in the midline (Fig. 1).

**Figure 1.**
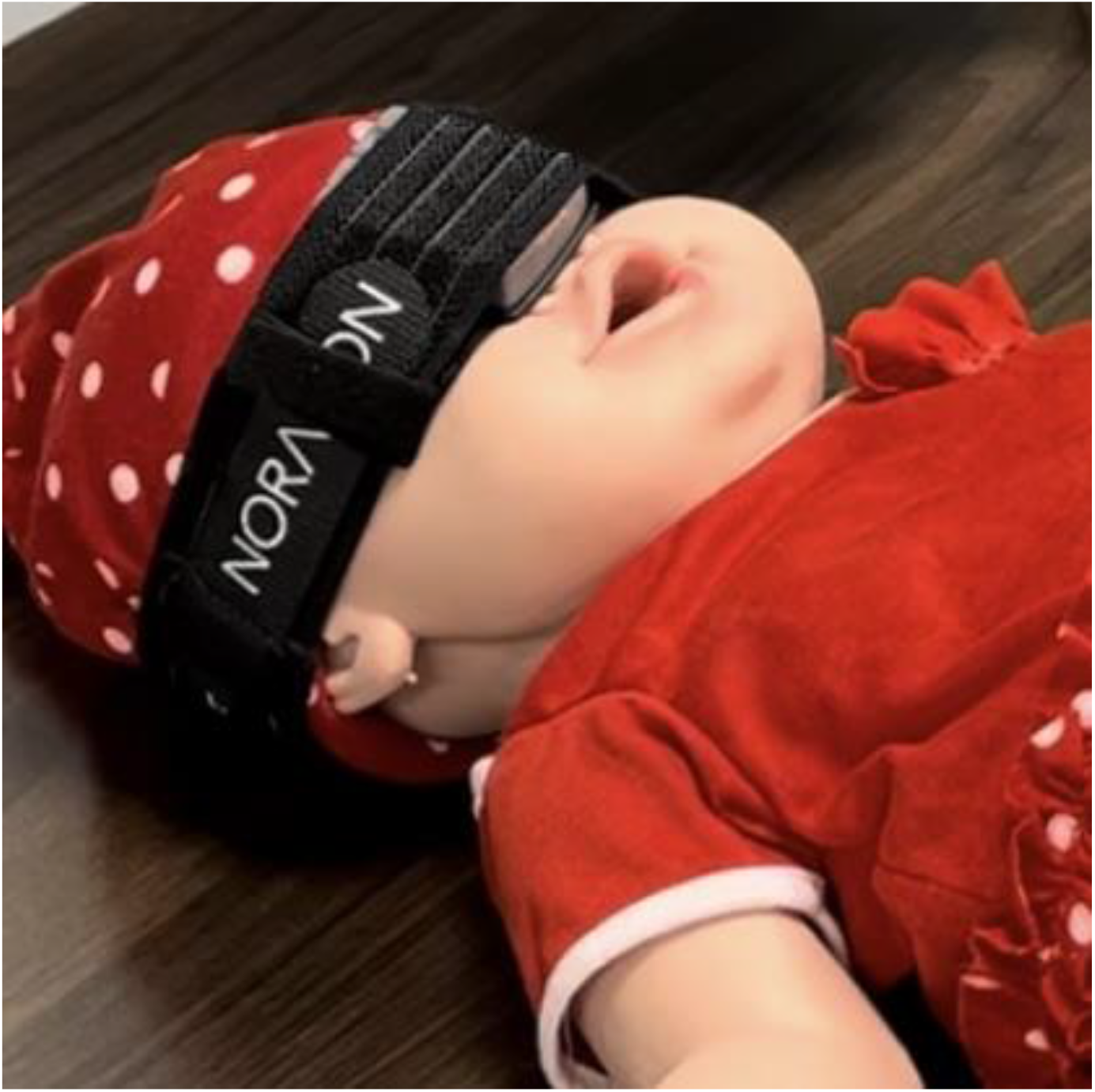
A photograph of the study design with an inertial measurement unit Velcroed around the silicone baby infant’s head to determine the head acceleration during a non-recommend handling and head impact with a table.

#### Procedures

Two participants (1M and 1F with no baby handling experience) were instructed to simulate placing an infant on a kitchen table (30 cm height and aluminum material). The participants were directed to hold the infant model by its torso, intentionally leaving the head unsupported, and then place the model supine on the table. Each participant was allowed five practice trials to familiarize themselves with the task and was explicitly instructed to avoid rough handling. Following this, five experimental trials were recorded for each participant.

Additionally, both participants were instructed to pick up the infant model, place their hands around the infant’s thorax, and shake the infant with maximum effort in a back-and-forth motion. This shaking task was performed once for approximately 5 seconds to simulate vigorous handling.

### Data Analysis

The inertial measurement unit was positioned on the infant’s head in an orientation such that X was vertical, Y was mediolateral, and Z was anteroposterior. Peak head acceleration was defined as the value of the first positive peak after the infant’s impact with the table. The peak vertical force was designated as the first spike immediately following head impact onto the table. The peak head acceleration for the Z-axis (anteroposterior) during the maximum effort shaking trials were extracted, rectified, and the highest 5 peak accelerations during the 5 second shake were selected and averaged. The peak head acceleration values for the Z-axis (anteroposterior) during the placing the infant on the table was calculated and averaged from the 5 trials.

To assess the magnitude of head acceleration between tasks, an independent t-test was conducted to compare the peak head accelerations during table placement with the peak head accelerations during the shaking with maximum effort of the infant model (SPSS, version 29, IBM Corp., Armonk, N.Y., USA). To assess inter-participant differences during the table placement task, an independent t-test was performed to compare the peak head accelerations during the table placement between the two participants’ five trials. To assess inter-participant differences during the maximum effort shake task, an independent t-test was performed to compare the peak head accelerations. Significance was determined at an alpha level of p < 0.05.

## Results

The summarized results are presented in Table 1 and 2. The average peak head acceleration of the infant models’ head when being placed on a table with the head unsupported was +30,952.67 ± 6,540.79 mg. In contrast, the peak head accelerations of the infant during a maximum exertion shake was significantly lower compared to the table condition (9,785.82 ± 3867 vs. 30,952.67 ± 6,540.79 mg, p < 0.0001, Fig. 3). There were no significant differences in peak head acceleration between the two participants during the table task (31,155.34 ± 9,404.02vs. 30,750 ± 2,778.71 mg, p = 0.93, Fig. 2). There were no significant differences in peak head acceleration between the two participants during the maximum effort shaking task (16,046.90 ± 9,539.06 vs. 12,289.10 ± 9,968.75 mg, p = 0.97).

**Table 1.** The summarized results of the mean peak head accelerations (mg) for each participant observed in placing the infant model on the table with the head unsupported and shaking the infant with maximum effort. There were no significant differences between individuals’ results in the maximum effort shaking and table placement task.

| Condition | Mean Peak Head Acceleration (mg) | Standard Deviation (mg) | Range of Head Acceleration (mg) | p-value |
| --- | --- | --- | --- | --- |
| Participant 1 (Table) | +31,155.34 | +9,404.02 | +19,234.40 - +43,406.30 | p = 0.93 |
| Participant 2 (Table) | +30,750.00 | +2,778.71 | +28,375.00 – 35,281.30 |  |
| Participant 1 Maximum Effort Shaking | +11,457.84 | +2,653.26 | -16,046.90 - +9,539.06 | p = 0.97 |
| Participant 2 Maximum Effort Shaking | +12,289.10 | +933.82 | -12,289.10 - +7,164.06 |  |

**Table 2.** The summarized results of the average peak head accelerations (mg) of both participants observed in placing the infant model on the table with the head unsupported and shaking the infant with maximum effort.

| Condition | Mean Peak Head Acceleration (mg) | Standard Deviation (mg) | Range of Head Acceleration (mg) | p-value |
| --- | --- | --- | --- | --- |
| Table Placement | +30,952.67 | +6,540.79 | +19,234.40 - +43,406.30 | p < 0.0001 |
| Maximum Effort Shaking | +11,430.48 | +1,875.42 | -16,046.90 - +9,539.06 |  |

**Figure 2.**
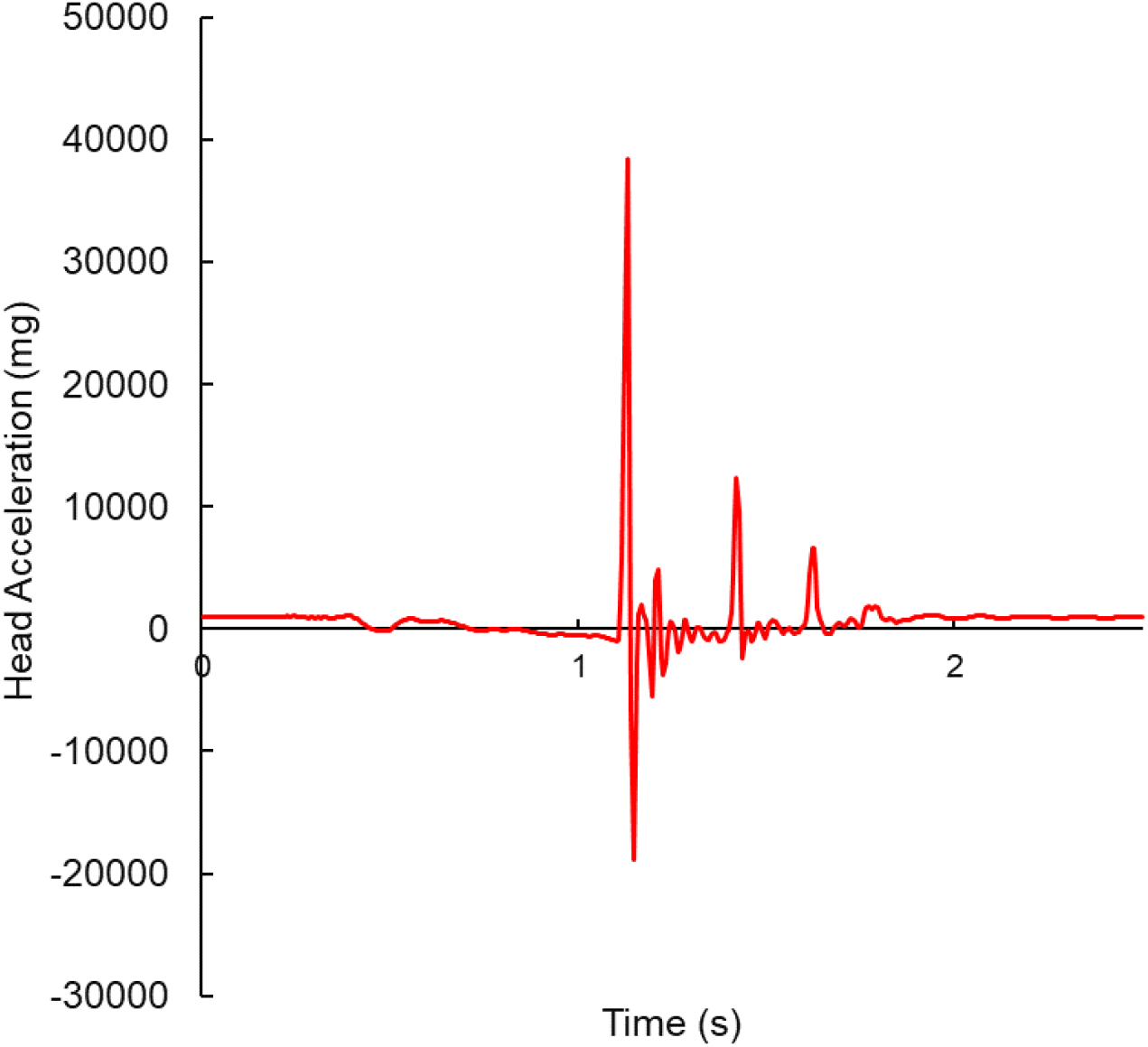
An example of a graph of the infant’s head acceleration (mg) during the impact with the table. The x-axis represents time in seconds.

**Figure 3.**
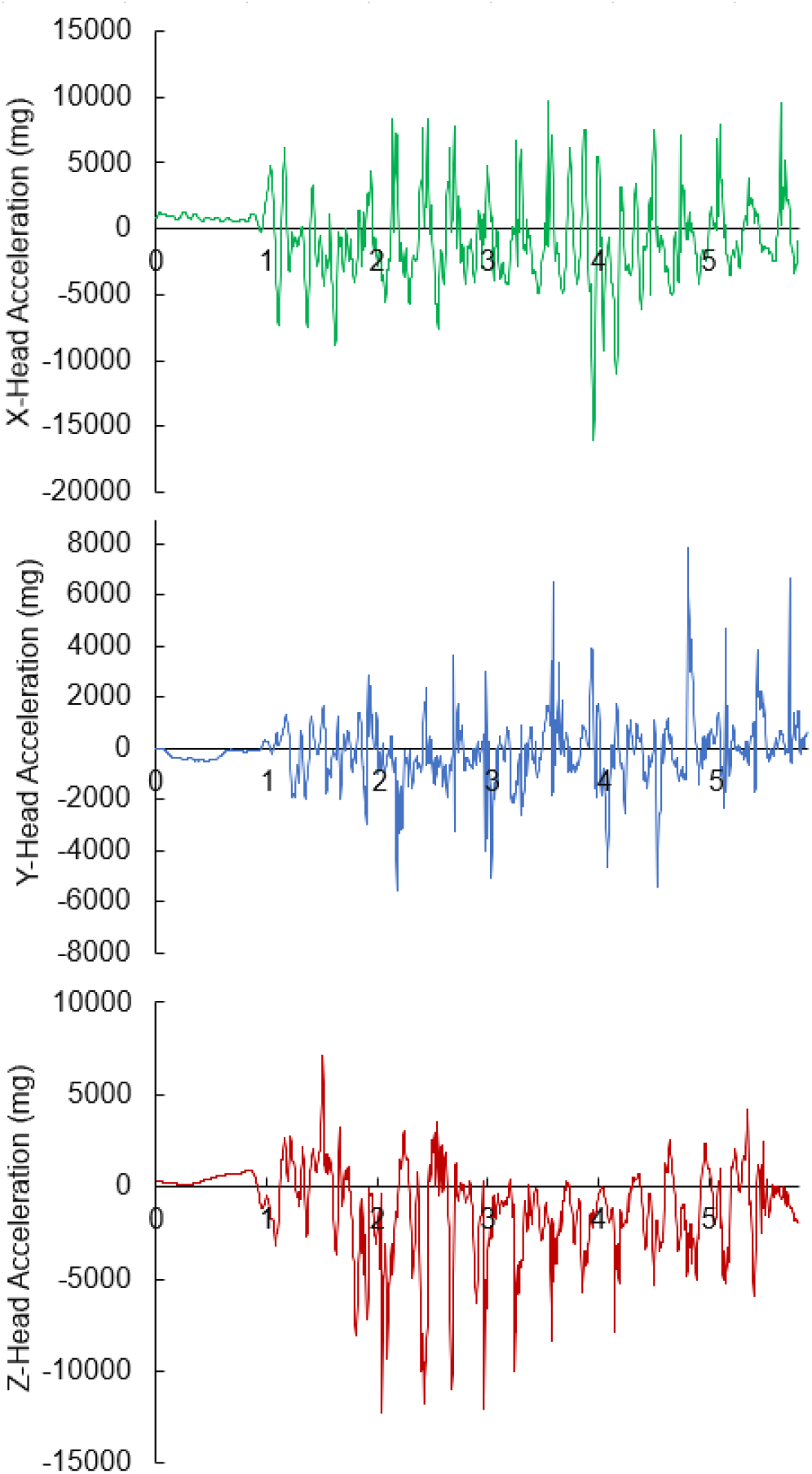
A graph of the head acceleration (mg) of the infant while being shaken at maximum-force. The x-axis represents the time in seconds.

## Discussion

Linear G-forces have been implicated in causing significant head injuries in infants, particularly in the context of abusive head trauma (AHT) [8]. In this study, we hypothesized that improper handling, such as placing an infant on a table without head support, could generate G-forces comparable to or even exceeding those associated with intentional shaking. Our findings confirmed this hypothesis, demonstrating that peak head accelerations during unsupported placement of the infant model on a table were over 300% higher than those recorded during maximum exertion shaking. These results highlight that G-forces produced in everyday scenarios, such as placing an infant on a hard surface without supporting the head, can exceed those generated during violent shaking.

When compared to known magnitudes, the G-forces observed in this study bring a new perspective in understanding the biomechanical forces on SBS. The peak head acceleration from placing an infant model on the table exceeds the 29.3 G-forces experienced in an 18-mph car crash and exceed the 5.2G-forces head accelerations recorded in a 5-mph bumper car collision [9]. The results from these prior studies, and compared to the results of this current study, highlight the potential for significant head injury in scenarios lacking malicious intent or due to improper handling of infants.

Previous studies have raised concerns about the traditional diagnostic criteria for SBS. From a biomechanical perspective, the G-forces induced on shaking a baby to create intracranial damage would induce significant spinal cord damage rather than head injuries [10,11]. This inherently makes sense as the infant’s head is the heaviest segment of the infant body, and the weak neck muscles would be unable to control forces acting on the vertebrae and spinal cord. This raises concerns about the role of other forms of trauma, including those that may occur unintentionally, in producing similar intracranial damage. While the autopsy report in this case study reported intramuscular cervical hemorrhages, it is important to note that the cervical hemorrhages aren’t as severe as expected from actual shaking of a baby [10]. It is plausible that an unsupported head while delivering back blows or placing the baby on the table with the head unsupported could have caused the intramuscular cervical hemorrhages given the high G-forces. Our findings suggest that it may be more appropriate to consider other forms of trauma, including those from unintentional mishandling, as contributing factors to head injuries in infants.The classic maximum exertion shaking was shown to produce 9-15 G-forces in prior work [5]. The prior study used an infant model that was 3.18 kg. This was not similar to the size of the 5.08 kg infant model used in this study. Even with the mass discrepancy, this study’s results confirmed prior work showing that maximum exerted shaking of an infant produced approximately 11 G-Forces [4,5]. The size of this model was smaller than the reported case. Since force is directly influenced by the mass of the object, a larger infant could experience even greater forces when placed on a table without head support. Furthermore, it is important to consider the improper handling of an infant as the head accelerations during the transfer of an infant to a table with the unsupported head resulted in 316% higher G-forces than maximum exertion of shaking an infant.

While the medical reports concluded that this was a case of infant homicide, it may be important to understand that blunt trauma may also stem from improper handling which can result in accidental trauma and high amounts of head acceleration in infants. Consideration of the entire medical incident is important to help determine whether the blunt trauma was accidental or intention. In this case study, the father’s account of the incident suggests unintentional harm may have occurred. This father found his infant in medical distress and possibly choking. The emotional stress and trauma of this on the parent could have caused him to handle his infant with the dire intention of reviving and seeking medical attention. The biomechanical data from this study demonstrated that such scenarios with no malicious intent can still pose a severe risk to an infant.

This study highlights the importance of supporting an infant’s head during handling to prevent accidental injury, and further highlights that SBS warrants additional context of the situation before ascribing malintent on a caregiver.

## Conclusion

This study offers insights into the biomechanical forces acting on an infant’s head during common caregiving scenarios, specifically when the head is not supported. The peak head accelerations observed during table placement without head support were significantly higher than those during maximum effort shaking, indicating that even non-malicious actions from improper handling can result in substantial g-forces. These findings emphasize the need for caregivers to be aware in supporting an infant’s head to prevent accidental injury. Additionally, our results highlight the importance of considering non-abusive causes when diagnosing SBS and suggest that further research is needed to explore the range of forces involved in everyday caregiving scenarios.

## Author Contributions

Conceptualization, J.L.C. and K.H.; Methodology, J.L.C. and K.H.; Software, J.L.C.; Validation, J.L.C., L.W., and K.H.; Formal Analysis, J.L.C.; Investigation, J.L.C. and L.W.; Resources, J.L.C.; Data Curation, J.L.C. and L.W.; Writing – Original Draft Preparation, J.L.C.; Writing – Review & Editing, J.L.C., L.W., and K.H.; Visualization, J.L.C. and K.H.; Supervision, J.L.C.; Project Administration, J.L.C and K.H..; Funding Acquisition, N/A

